# Genome-wide analysis of dry (Tamar) Date palm fruit color

**DOI:** 10.1101/2022.12.12.520041

**Authors:** Shameem Younuskunju, Yasmin A. Mohamoud, Lisa Sara Mathew, Klaus F. X. Mayer, Karsten Suhre, Joel A. Malek

## Abstract

Date palm (Phoenix dactylifera) fruit are an economically and culturally significant crop in the Middle East and North Africa. There are hundreds of different commercial cultivars producing dates with distinctive shapes, colors, and sizes. Genetic studies of some Date palm traits have been performed, including for date palm sex-determination, sugar content and fresh fruit colour. In this study, we used genome sequences and image data of 199 dry date fruit (Tamar) samples collected from 14 countries to identify genetic loci associated with the color of this fruit stage. Here, we find loci across multiple linkage groups (LG) associated with dry fruit color phenotype. We recover the previously identified VIR genotype associated with fresh fruit yellow or red color and new associations with the lightness and darkness of dry fruit. This study will add resolution to our understanding of the date palm fruit color phenotype especially at the most commercially important tamar stage.

## Introduction

Date palm (*Phoenix dactylifera*) is one of the oldest and most economically important fruit crops in the Middle East and North Africa (Chao & Krueger, 2007; Weiss, Zohary, & Hopf, 2012). While there are thousands of cultivars or varieties, likely, only a few hundred are commercially important (Zaid & Arias-Jimenez, 1999). These cultivars produce fruit (Dates) that have distinctive shapes, colors, and sizes of fruit. Fruit development and ripening involve many complex biological processes, and color changes of the fruit are closely associated with the ripening stage (Abbas & Ibrahim, 1998). Dates have five different development stages: Hababauk, Kimri, Khalal, Rutab, and Tamar (Al-Mssallem et al., 2013; Siddiq & Greiby, 2013). Hababauk and Kimri are the first two development stages, and fruit skin color is whitish-green. In the Khalal stage, dates partially ripen and gain maximum size and weight. During this stage, fruit color changes from green to yellow or red depending on the cultivar. Dates fully mature in the Rutab stage and the color begins its change to brown. Tamar is the final stage of ripening, during which fruit water content is reduced to less than 25%, sugar content increases to 70 to 80 % and the color turns dark brown. During fruit ripening process amino acids act as building blocks for the synthesis of key intermediates and end products (Seymour, Taylor, & Tucker, 2012) and the enrichment of both free amino acids and color or flavor-conferring phenylpropanoids was observed in the early ripening stage of dates as in other fruits (Diboun et al., 2015).

Dates are classified as climacteric fruit like other commercial fruit such as apples, bananas and peach where ethylene functions as a key regulator in fruit ripening (Abbas & Ibrahim, 1998; Al-Qurashi & Awad, 2011; Serrano, Pretel, Botella, & Amoros, 2001). Fruit ripening is associated with an increase in respiration rate, burst of ethylene production, oxidation process, sugar accumulation, chlorophyll break down, pigment synthesis and other processes (Barreveld, 1993; Osorio, Scossa, & Fernie, 2013; Stepanova & Alonso, 2005). Anthocyanin is a class of secondary metabolite synthesized in higher plants and plays a crucial role in pigmentation (red, pin Later k, purple & blue) in fruit and vegetables (Kong et al., 2003; Kayesh E et al.,2013). In anthocyanin biosynthesis, the R2R3 MYB transcription factors act as a regulator (Xie et al., 2020). Previous studies from Hazzouri *et al*. (K. M. Hazzouri et al., 2015; Khaled M. Hazzouri et al., 2019) revealed that a retrotransposon insertion (named Ibn Majid) and a start codon mutation in an R2R3 MYB transcription factor encoded by the ortholog of the oil palm VIRESCENS gene (VIR gene), are likely the key genetic changes leading to the red and yellow color phenotype in dates at the Khalal stage (fresh fruit). Studies by Awad (2007) and Shareef (2020) showed that the color transition form from yellow to brown (Khalal-to-Rutab) is associated with an increase in endogenous Abscisic Acid (ABA) concentration (Awad, 2007; Shareef & Al-Khayri, 2020). A recent study by Saar Elbar *et al*. provided evidence of gradual increase of pulp ABA during the ripening stage (Khalal-to-Rutab-to-Tamar) (Elbar et al., 2022). They also showed that the color transition from yellow to brown (Khalal-to-Rutab) was preceded by an arrest of xylem-mediated water transport into water.

Studies of the genetic basis of Date palm’ traits have significantly increased due to the crop’s economic importance. Genome-wide association studies (GWAS) are a powerful method for mapping the association between genetic variation and phenotype (Cantor, Lange, & Sinsheimer, 2010; Korte & Farlow, 2013). Previous GWAS in date palm identified significant associations for fruit sugar content (Khaled M. Hazzouri et al., 2019; Malek et al., 2020), confirmation of previous findings of the sex determination region on LG12 (Khaled M. Hazzouri et al., 2019; Lisa S. Mathew et al., 2014) and fresh fruit color (Khaled M. Hazzouri et al., 2019). A goal of GWAS is to identify the association between the variance of the phenotype of interest and genomic region or loci at genome-wide significance. Challenges in conducting GWAS in date palm exist including the use of clonal propagation, rare existence of outbred populations and geographical population structure. The population structure and cryptic relatedness have the potential to confound the GWAS results and can lead to false discoveries (Chen et al., 2016; Horton et al., 2012; Vilhjálmsson & Nordborg, 2013). Along these lines, multiple studies (Chaluvadi, Khanam, Aly, & Bennetzen, 2014; Flowers et al., 2019; Zehdi-Azouzi et al., 2015), including our own (L. S. Mathew et al., 2015) have confirmed two major subpopulations (western and Eastern populations) in the date palm population. Indeed, our recent study reported that at least three and possibly four novel subpopulations contribute to the current date palm population (Mohamoud et al., 2019) making population structure an important consideration in any date palm genome-wide association study. Different algorithms have been developed to correct for population structure and increase the computational efficiency and statistical power in GWAS (H. M. Kang et al., 2008; Q. Wang, Tian, Pan, Buckler, & Zhang, 2014; Y. Zhang et al., 2018). Fixed and random model Circulating Probability Unification (FarmCPU) is a statistically powerful and computationally efficient GWAS method to control spurious associations (Liu, Huang, Fan, Buckler, & Zhang, 2016; J. Wang & Zhang, 2021). The iterative usage of the fixed effect and random effect models in the FarmCPU method incorporates population structure and kinship matrix as covariates and eliminates the false positive and false negative association results. We therefore hoped to implement these methods in the challenge of GWAS in the highly structured date palm population.

In this study, we conducted GWAS using 199 date palm samples to identify the significantly associated genetic loci and possible candidate genes with the color variation of Tamar stage fruit (dry fruit). The samples used in this study are extensively diverse in their country of origin and variety, collected from 14 countries.

## Materials and Methods

### Sample collection and genome sequencing

The Qatar date fruit biobank is a collection of date fruit samples from across the date palm growing region, including Qatar, United Arab Emirates (UAE), Iran, Saudi Arabia, Egypt, Pakistan, Libya, Tunisia, United States of America (USA), Morocco, Jordan, Sudan, Oman and Spain (N Stephan et al. 2018). We used 199 date fruit samples from the collection, attempting to include the most important commercial cultivars as well as some lesser-known varieties (Supplementary file 1). DNA from fruit was extracted, sequencing libraries were constructed from total DNA (Mathew LS et al., 2015) and sequenced on the Illumina HiSeq 2500/4000 instruments using 150 bp length reads. All sequencing steps were done as described in Thareja et al (Thareja et al., 2018).

### Phenotypic data

Photographs were taken of 5 to 11 Tamar stage fruit from each cultivar along with a color-checker card and a ruler as a size standard (Supplementary Figure 1). Pictures were taken on a white background with a digital camera. For more detailed description please refer to Stephan et al (Stephan et al., 2018). Out of 199 samples, only 179 fruit samples were available for phenotypic data and used for the association study. We measured fruit color by recording the intensity of the Red (R), Green (G) & Blue (B) color channels (RGB) in each photograph for each fruit using MIPAR Software (Sosa et al.,2014). The color channel intensity of multiple fruit was calculated separately and then averaged for each cultivar. The average intensity of each color channel (R, G &B) and the ratio of the color channel such as R/G, R/B & G/B were used as phenotype data for the association mapping study.

### Genome alignment and SNP calling

Raw Illumina reads of 199 samples were processed to remove adaptor sequences using Trimmomatic software (v0.39). We used the ‘ILLUMINACLIP:2:30:10 MINLEN:50’ settings to keep the read pairs with a minimum length of 50bp for further processing. Adaptor removed reads were aligned to the PDK50 reference genome (PRJNA40349) using bwa mem software (v0.7.15-r1140) (Heng Li, 2013) with default parameters. Reads with supplementary (split/short) alignments were marked by the bwa mem -M option. Each sample alignment was validated with ‘ValidateSam’ (Picard) (http://broadinstitute.github.io/picard) to ensure no errors in output BAM files. Before variant calling, duplicate reads were marked using GATK MarkDuplicates (gatk-4.1.7.0) (DePristo et al., 2011). The following GATK function, ‘FixMateInformation’ and ‘SetNmMdAndUqTags,’ were used to ensure consistent paired-read information and fix the NM, MD, and UQ tags in the sorted BAM files. Variant calling was performed using GATK HaplotypeCaller with GVCF mode. Reads with mapping quality less than 20 and duplicate marked reads were excluded from the variant call. Hard filter parameters were applied to filter raw SNPs and INDELS using bcftools (v1.10.2) (H. Li, 2011). The raw SNPs were filtered using ‘QD<2.0 || FS>60.0 || MQ<40 || SOR>4.0 || MQRankSum<-10 || MQRankSum>10 || ReadPosRankSum<-6’ parameters, and INDELS were filtered with ‘QD < 2.0 || FS > 200.0 || ReadPosRankSum <−20.0’ parameters. Thereafter we utilized QC filter on SNP data. We excluded samples with greater than 70% missingness and SNPs with depth > 11000 summed across the samples in QC analysis. Genotypes were marked as missing if DP was below 10, genotype call rate less than 80%, or a minor allele frequency below 0.01. Finally, multiallelic SNP (-max-alleles 2) and SNPs with Hardy–Weinberg Equilibrium p-value less than 1e-06 (--hwe 1e-06) were filtered out using Vcftools (V0.1.16). The resulting samples and SNPs were used for further analysis.

### Linkage Disequilibrium (LD) Analysis

Linkage Disequilibrium (LD) was calculated using PopLDdecay software (C. Zhang, Dong, Xu, He, & Yang, 2018) with -MaxDist 200 -MAF 0.05 option. The pairwise correlation coefficient (r^2^) was calculated for all SNPs in 200kb window with minor allele frequency greater than 5% for the whole genome and 18 linkage groups separately (LG). The decay of LD was calculated by plotting pairwise r^2^ onto genetic distance between the SNP pairs using an approach mentioned in Marroni *et al*. (Marroni et al., 2011) and then calculated the distance at which pairwise correlation coefficient (r^2^) is half its maximum value as half decay distance.

### Genome-wide association study

GAPIT (v3) R package implemented FarmCPU (J. Wang & Zhang, 2021) method was used for the genome-wide association to fruit color phenotype. We ran LD pruning on QC passing SNP data to improve the computational efficiency of the GWAS analysis. LD pruning was executed with plink software (Purcell et al., 2007) using --indep-pairwise 500 50 0.99 option. We used the first four principal components (PCA) and Kinship matrix as covariates to correct population structure. PCA and Kinship matrix was generated by GAPIT using LD pruned SNPs data set. VanRaden algorithm in GAPIT R package used for calculating kinship. In this study, an FDR-adjusted p-value of 0.05 (5%) was used as a cut-off value for identifying the list of significant SNPs associated with date palm fruit’s color phenotype. QQ-plot and Manhattan plots were generated from the GWAS result using the R package CMPlot.

### VIR genotyping and significant SNP effects on Tamar stage fruit color

The genomic region of the VIRESCENS gene (VIR gene) and Ibn Majid retrotransposon from the Barhee BC4 genome (Hazzouri *et al*. study,2019) were aligned against the PDK50 reference genome, and the coordinates of its insertion and termination codon in the VIR gene noted. Following the same procedure used in Hazzouri *et al*., we manually inspected all the QC filtered samples to identify the VIR genotypes (VIR^+^/VIR^+^, VIR^+^/VIR^IM,^ and VIR^IM^/VIR^IM^) using JBrowse2 software. In this study, samples were classified into two groups (i): homozygous VIR^+^/VIR^+^ genotype (ii): homozygous VIR^IM^/VIR^IM^ genotype. That is, only homozygous genotypes were used for further analysis to avoid confusion about the effect of the heterozygous genotypes. To investigate the effect of loci on dry fruit color identified in this study, analysis of our SNPs was conducted within each homozygous VIR sample group.

### Candidate gene and variant analysis

±150kb region on both sides of significant SNPs (total 300kb) were considered as potential regions for possible candidate genes and variant analysis. Possible gene sequences and coordinates were mapped from the potential region using the GFF3 gene annotation file (PDK50.gff3) of the PDK50 reference genome. Gene ontology of mapped gene sequences was performed using Blast2Go software (Conesa & Götz, 2008). Literature search and Blast2Go results were used to identify the function of the candidate genes. All SNPs and INDELs from the potential regions were annotated using SNPeff software. LD R^2 value of each SNP from the candidate region was calculated against the significant SNPs from the GWAS result using PLINK software. Amino acid substitution effects on the protein function of the candidate SNPs were analysed using SIFT (Sim et al., 2012).

### Structural variation Analysis

Clipped, discordant, unmapped, and indel reads from each sample were used for structural variation analysis. The reads were collected from samples having GWAS significant SNPs with homozygous allele alternative (ALT) to the reference genome. Reads from each sample were grouped based on homozygous ALT alleles and assembled separately using spades software (SPAdes-3.15.2) (Bankevich et al., 2012). According to the manual recommended parameters, the mapping and annotation of structural variation from assembled contigs were performed using Assemblytics software (Nattestad & Schatz, 2016). Assembled contigs were aligned against the reference genome using nucmer software (Marçais et al., 2018). We then mapped structural variation breakpoints and annotated them using the Assemblytics software.

### RNA seq Analysis

RNA-seq expression analysis was performed using data from Hazzouri *et al*.*’s* study (Khaled M. Hazzouri et al., 2019). We used RNA-seq data of three or four replicates of varying days post-pollination (45,75,105,120 & 135 days) in Kenezi (dark brown color fruit) and Khalas (light brown color fruit) fruit. RNA-seq fastq files were downloaded from SRA (PRJNA505138). The reads were aligned to the reference genome using STAR split read aligner (V 2.7.1a) (Dobin et al., 2013) with two pass alignment. Thereafter, reads count per gene were calculated using the featureCounts program with the following parameter ‘-p --countReadPairs -t exon -g gene_id -s 0’. Taking DESeq2 with the median-of-ratio method, read normalization was performed for the expression study.

## Results

### Phenotypic data

In the Tamar stage, among other changes, date palm fruit color varies from light brown to dark brown. For the genome-wide analysis study, we used the intensity of Red, Green & blue color channels and the ratio of Red/Blue (R/B), Red/Green (R/G), and Green/Blue (G/B) as phenotype. Manual analysis of fruit images with the ratio of Red and Blue channel intensity (R/B) showed that higher values of R/B are associated with light brown and lower R/B values with dark brown (**Figure 1**). We, therefore, focused our analysis on R/B color phenotype for the genome-wide analysis of fruit color.

**Figure 1.**
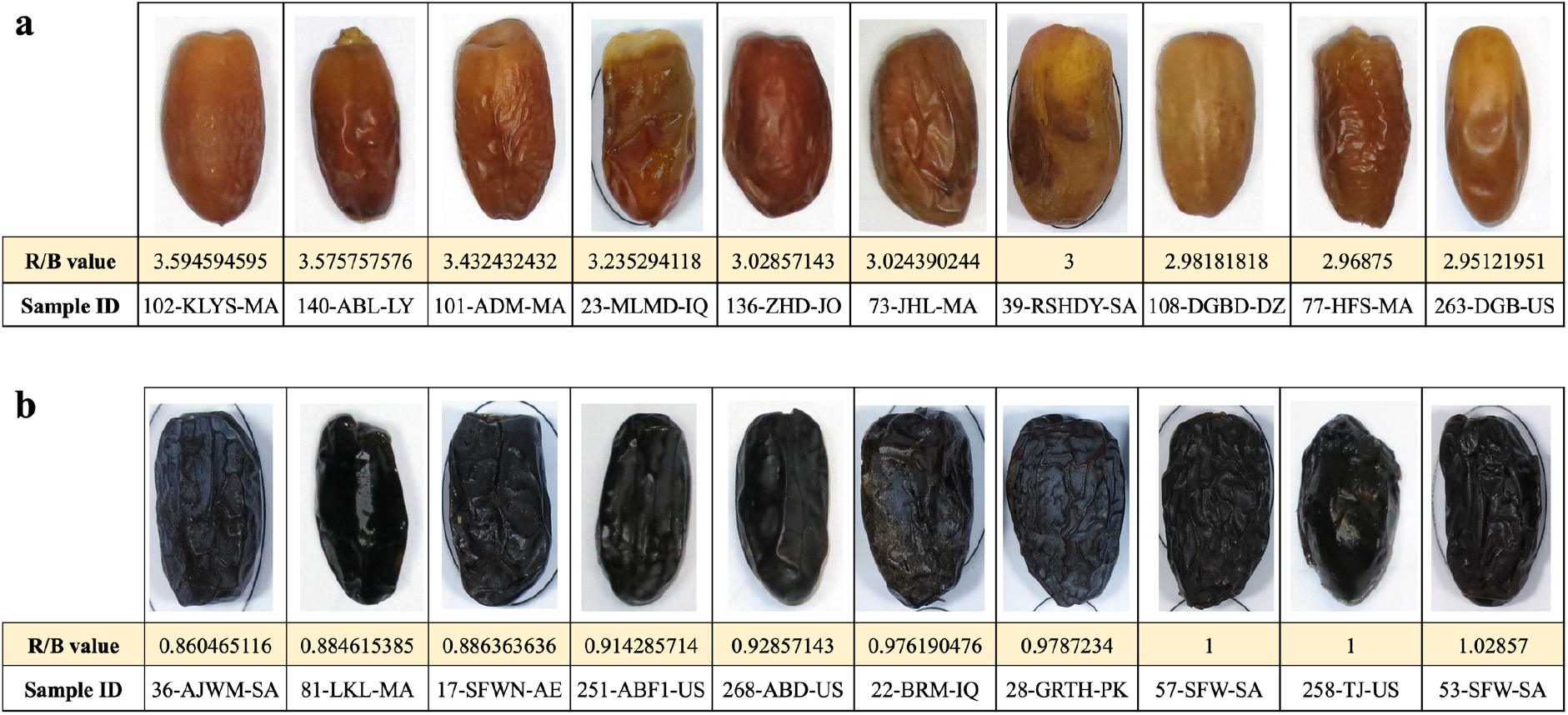
Comparison of Fruit images with R/B phenotype value. Fruit color ranges from dark to light brown color as R/B value increases. The maximum R/B value among 179 samples is 3.59, and the minimum value is 0.86. **a:** Fruit images of 10 samples which the highest R/B value. **b:** fruit images of 10 samples which the lowest R/B value..

### LD analysis

Our GWAS mapping population (n=188) contain 68 samples of the eastern cultivar (36 %) and 106 samples of the western cultivar (56.8%). LD analysis was performed for the whole genome and each Linkage group of GWAS population (n=188) and separate for the whole genome of eastern and western cultivars. The pairwise correlation coefficient (r^2^) for the whole genome of the GWAS population decreased to half of its maximum value at 25.9 kb (Figure 2a), which is close to the LD decay distance (22.9 kb) found by Hazzouri et al. (Khaled M. Hazzouri et al., 2019). The LD decay was slower for LG12 and LG16. The r^2^ decreased to half of its maximum value at 38.4 kb and 31.9 kb. The LD decay was faster for LG 4, LG 7 and LG 15. The r^2^ decreased to half of its maximum value at 18.9 kb, 20.5 kb and 17.2 kb respectively (Table 1, Supplementary Figure 2). The LD decay analysis of the eastern and western cultivars shows that r^2^ decreased to half of its maximum value at 21.73 kb and 35.85 kb respectively (Supplementary Figure 3).

**Figure 2.**
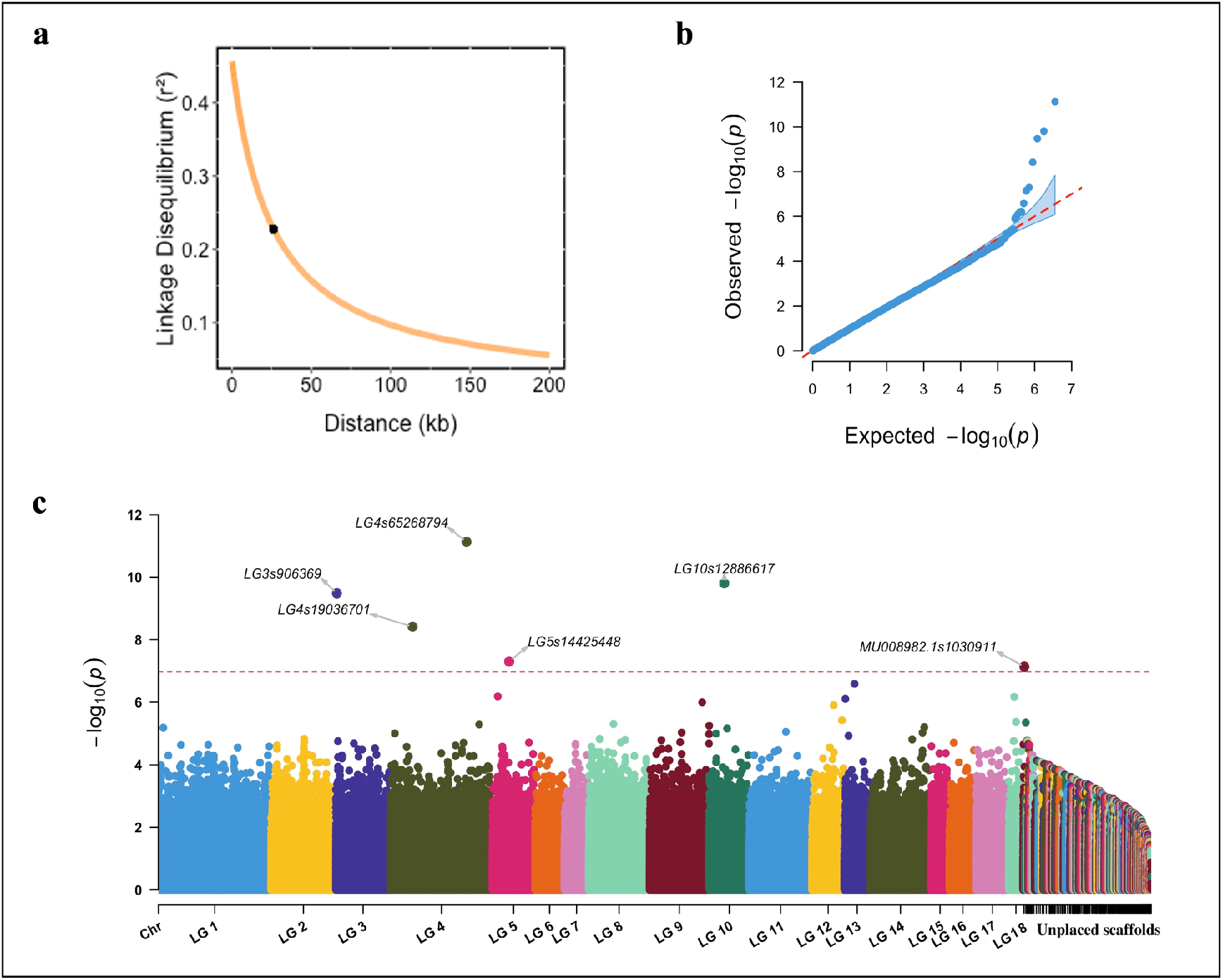
GWAS analysis of tamar stage fruit color (R/B) of date palm. **a**. decay of linkage disequilibrium with physical distance in the GWAS mapping population. The half decay distance is 25.9 kb. **b & c :** GWAS of fruits date palm color(n=188), (b): QQ plot (b) and (c) Manhattan plot using the LD pruned SNPs set (3,541,727 SNPs) for all linkage groups (LG) and unplaced scaffolds.

**Table 1:** Decay of Linkage Disequilibrium (LD) with physical distance in each linkage group (LG)

| LG id | half LD decay ($r^2$) | half LD decay distance (kb) |
| --- | --- | --- |
| LG1 | 0.2282535 | 27.94 |
| LG2 | 0.2282521 | 23.81 |
| LG3 | 0.2282519 | 23.53 |
| LG4 | 0.2282495 | 18.90 |
| LG5 | 0.2282531 | 26.51 |
| LG6 | 0.2282532 | 26.92 |
| LG7 | 0.2282505 | 20.58 |
| LG8 | 0.2282518 | 23.14 |
| LG9 | 0.2282527 | 25.48 |
| LG10 | 0.2282519 | 23.36 |
| LG11 | 0.2282527 | 25.47 |
| LG12 | 0.2282558 | 38.49 |
| LG13 | 0.2282509 | 21.26 |
| LG14 | 0.2282532 | 27.05 |
| LG15 | 0.2282484 | 17.27 |
| LG16 | 0.2282545 | 31.90 |
| LG17 | 0.2282533 | 27.16 |
| LG18 | 0.2282515 | 22.44 |

### Association study of Tamar stage fruit color

Genotyping of date palm samples using GATK HaplotypeCaller produced a filtered call set of 31,373,292 SNPs. Following the application of quality control filters (QC filters), we identified 10,183,834 SNPs across 188 samples. The association study was performed for all fruit color phenotypes such as Red, Green, Blue, R/B, R/G & G/B using the FarmCPU GWAS method. For computational efficiency of FarmCPU, we chose to remove LD-correlated SNPs from the GWAS panel. LD pruning using PLINK resulted in 3,541,727 SNPs. GWAS results of all phenotypes except the ‘Blue’ color channel phenotype showed overlapping significant SNPs. After manual analysis, we chose to focus on the GWAS results of the R/B phenotype for the reasons mentioned above. Q-Q plot of FarmCPU genome-wide association results with the R/B phenotype shows a sharp deviation from the expected P-value distribution in the tail area and lamda score as 1.02 (Figure 2b) showing good control of both false positives and false negatives. The FDR adjusted P-value cut-off of the R/B GWAS result shows that 6 SNPs are significantly associated with the phenotype (Table 2 and Figure 2c). These SNPs are from multiple linkage groups (LG) such as LG3, LG4, LG5, LG10 and one unplaced scaffold MU008982.1. A SNP from LG4 (LG4s65268794) has a highly significant p-value (10e-12), and it is common between GWAS results of the Red (R), R/B and R/G phenotypes (Figure 2c, Supplementary Figure 4 & 7). SNP LG10s12886617 from LG10 is common in R/B and G/B phenotypes (Supplementary Figure 8). The association result of R/G also shows a significant SNP in LG10 (LG10s12504803), which is very close to LG10s12886617 SNP. The significant SNPs from the association result of R/B phenotypes were used for identifying the loci and possible candidate genes associated with the color phenotype of Tamar stage fruit.

**Table 2:** List of significant SNPs associated with R/B phenotype from FarmCPU GWAS result. An FDR-adjusted p-value of 0.05 (5%) was used as a cut-off value for identifying the list of significant SNPs associated with the phenotype

| SNP | LG | Position | P.value | FDR<br>P.values | maf |
| --- | --- | --- | --- | --- | --- |
| LG4s65268794 | LG4 | 65268794 | 7.40E-12 | 2.62E-05 | 0.41620112 |
| LG10s12886617 | LG10 | 12886617 | 1.56E-10 | 0.000276 | 0.19273743 |
| LG3s906369 | LG3 | 906369 | 3.29E-10 | 0.000388 | 0.29329609 |
| LG4s19036701 | LG4 | 19036701 | 3.84E-09 | 0.003401 | 0.17877095 |
| LG5s14425448 | LG5 | 14425448 | 4.98E-08 | 0.035277 | 0.08938547 |
| MU008982.1s1030911 | MU008982.1 | 1030911 | 7.25E-08 | 0.042803 | 0.20391061 |

### Significant SNP’s genotypic effects on the Tamar stage fruit color

We conducted further analysis of the SNPs identified here as associated with fruit color (Table 2). Visual inspections, shows that allelic variation in these SNPs are indeed associated with light (high R/B value) or dark brown color (low R/B value) fruit (Figure 3a and Supplementary Figure 9). In our results, the R2R3 transcription factor gene (ortholog of the oil palm VIRESCENS gene -VIR gene), previously reported to regulate red or yellow fruit color at khalal stage (fresh fruit) (K. M. Hazzouri et al., 2015; Khaled M. Hazzouri et al., 2019), is present 16kb away from the SNP LG4s65268794 on LG4. As validation, we separated our samples by the VIR genotype and compared them to their fruit color. Our results confirm that samples were classified into light brown, dark brown and mixed color of light & dark fruit based on the VIR genotype (Supplementary Figure 10). Out of 188 samples, 27 fruits sample are homozygous wild type (VIR^+^/VIR^+^) and show dark brown color fruit, while 61 are homozygous for the transposon insertion (VIR^IM^/VIR^IM^) and show light brown color fruit. 100 have VIR^+^/VIR^IM^ genotypes that show the mixed color of the fruit (light & dark brown) in which VIRIM act as dominant or semi-dominant by interfering with expression of the wild type VIR^+^ allele. We also identified start codon mutation (ATG to ATA) of R2R3 transcription factor gene (called as VIR^saf^) in homozygous wild-type fruit, previously reported in Hazzouri et al’s study (2019). VIR^+^ is haplosufficient in VIR^+^/VIR^saf^ genotype and it produces dark brown color fruit like homozygous wild type (Khaled M. Hazzouri et al., 2019).

**Figure 3.**
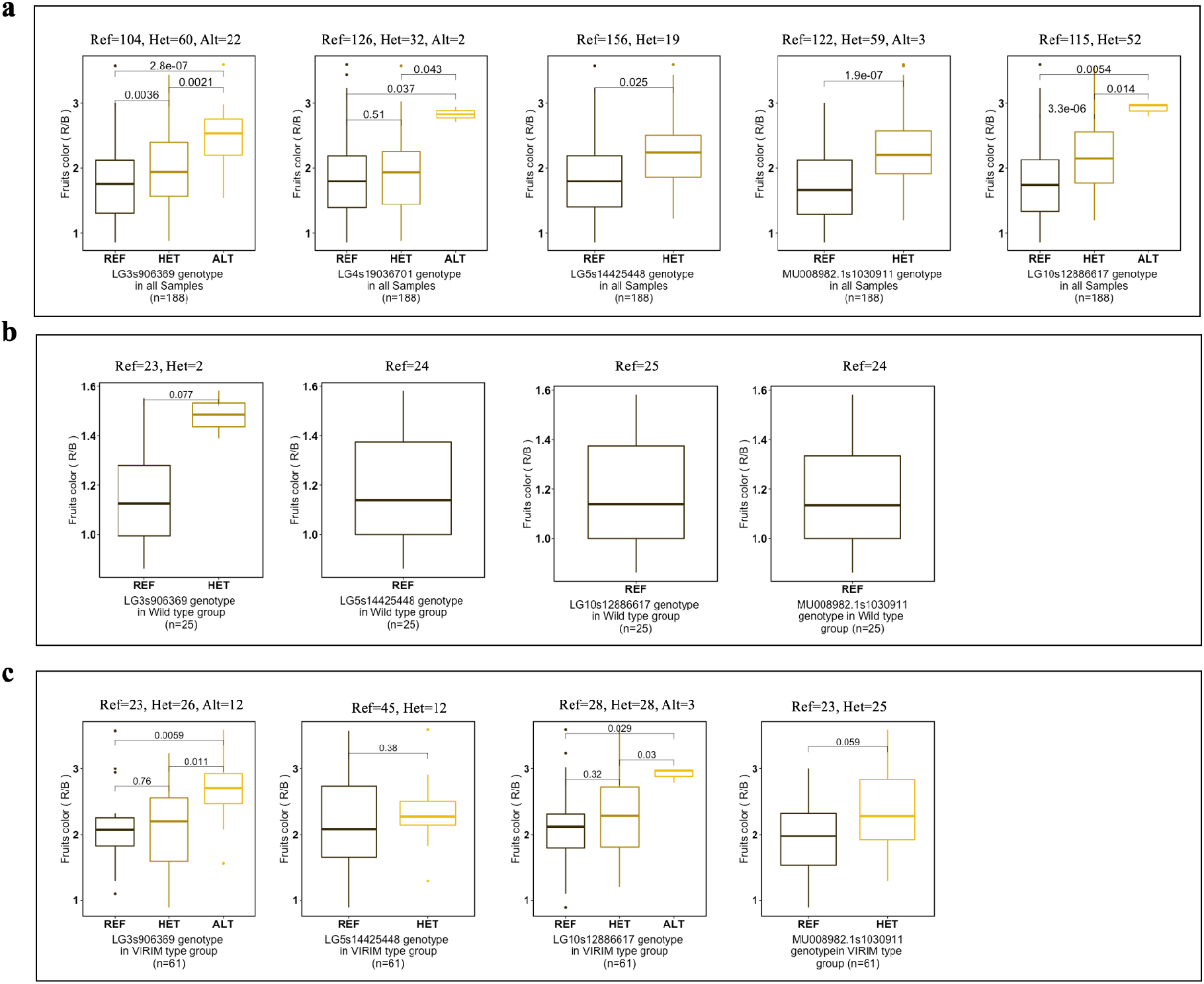
Boxplot distributions analysis of fruit color phenotype (R/B) by genotypes of significant GWAS SNPs from this study. Samples were classified into three group: (**a**) all 188 samples group, (**b**) wild type VIR group: samples with VIR^+^/VIR^+^ or VIR^+^/VIR^saf^ (n=25) and (**c**) VIRIM group: samples with VIR^IM^/VIR^IM^ genotype (n=61) and then further separated by genotypes of significant SNPs identified in this study. P-values were calculated using by Wilcoxon statistical test. The X-axis represents the SNP’s genotypes, and Y-axis represents R/B phenotypic value. Results show significant association with fruit color even when samples are separated by the VIR genotypes associated with red/yellow fresh fruit color.

For further understanding of color variation of dry fruit beyond the R2R3 transcription factor, we analyzed fruit color phenotype by the genotype of significant SNPs from our GWAS (Table 2) while excluding the LG4s65268794 SNP that corresponds to the genotype state of the VIR gene. The analysis was conducted in all 188 samples and then in groups separated by VIR genotype (Figure 3a, Supplementary Figure 11). Genotypes of SNP LG4s19036701 could not distinguish the fruit color in the VIRIM sample group, likely because of linkage on the same chromosome as the VIR gene, and so was excluded from this specific analysis. The results from analyses of the combined 188 samples showed significant associations (Wilcoxon test) with the color phenotype as expected (Figure 3a). To confirm the genotypic effect of these SNPs within the genetic background of the VIR genotype sample group, we performed the phenotype and genotype analysis in each group separately. We observed that even within the various VIR genotype groups that are key to red and yellow fresh fruit color, the SNPs identified here often further distinguish fruit color significantly (Figure 3b & 3c) suggesting these regions may play a role in dry fruit color or a more fine-grain role in fresh fruit color that was previously undetected. The result from the homozygous VIR wild-type fruit group, excluding the VIR^saf^/VIR^saf^ genotype samples, (total 25 sample) shows that fruit’s darkness decreases (R/B value increases) when the sample is heterozygous for SNP LG3s906369 while darkness increases (R/B value decreases) when the sample is homozygous for the reference allele (Figure 3b). SNP LG5s14425448, LG10s12886617 and MU008982.1s1030911 have only homozygous REF alleles in the group, so did not the numbers required to show any fruit color variation. The VIRIM sample group shows that the lightness of the fruit color increases (R/B value increases) when the sample is homozygous ALT or HET allele of SNP LG3s906369, LG5s14425448, LG10s12886617 and MU008982.1s1030911 (Figure 3c). Results from both groups agree with the 188-sample group result; fruit color can be distinguished based on the genotypic variation of the SNPs. Manual fruit color analysis by the genotypes of SNP LG3s906369 on VIRIM sample group agrees with this result (Figure 4). Sample group details and genotype information of SNPs are mentioned in the Supplementary file 2.

**Figure 4.**
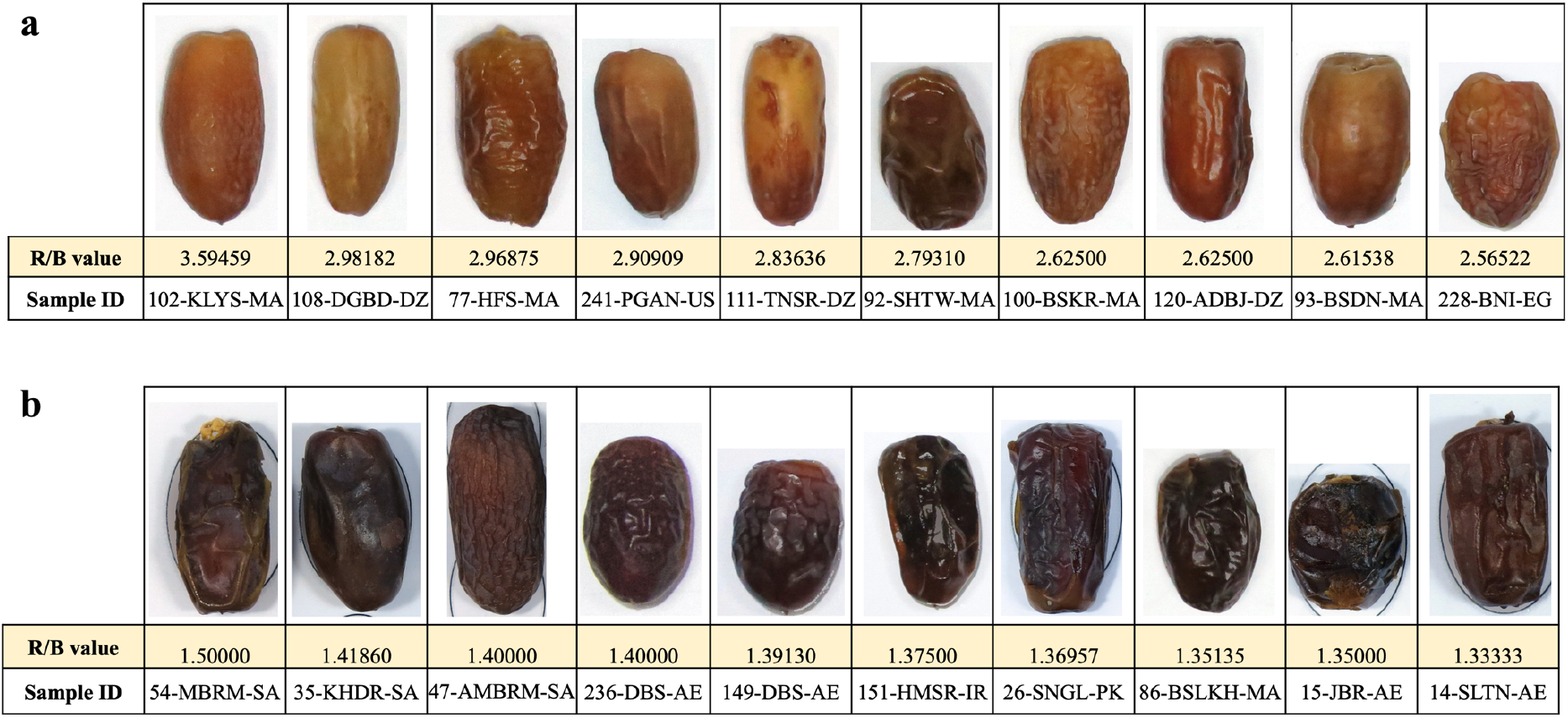
Fruit color differences within the homozygous VIRIM sample group (yellow fresh fruit color) when categorized by the genotypes of SNP LG3s906369. Fruits were grouped together which are homozygous for the **(a)** ALT or **(b)** REF allele of SNP LG3s906369. Results shows that the lightness of the color increases (R/B value increases) when the sample is homozygous ALT allele of SNP LG3s906369.

### Candidate genes and SNPs annotation

We considered potential candidate genes when they were within ±150kb of a significantly associated SNP (total of 300kb). While this is broad given average LD decay, we wanted to ensure the possibility of capturing linked genes that may be outside the standard LD decay size. There were 117 genes present across all the potential regions of significant SNPs from the GWAS result of the R/B color phenotype (Supplementary file 3). The putative gene functions were assigned by similarity using blast2Go software. The blast2Go and literature search results show that many genes related to fruit ripening and pigmentation are present around the region of significant SNPs (Table 3) and many are present within a 50kb of the significant SNPs. The gene expression analysis using the RNA-seq data for Kenezi (dark brown color) and Khalas (light brown color) fruit varieties shows that many genes from the potential candidate regions were expressed in fruit during the various days post-pollination (dpp) (Figure 5). R2R3 transcription factor gene from LG4 was expressed late in the development stage in both dark and light brown color fruit cultivar (dpp 105, 120 and 135). The R2R3 transcription factor gene has reduced expression in light color fruit variety compared with dark. RING/U-box superfamily protein from LG10 was expressed at dpp 105 to 120 in the light fruit variety (peaking at dpp 120) compared to the dark color fruit. Other genes such as Protochlorophyllide reductase (LG3), Basic helix-loop-helix (BHLH) DNA-binding superfamily (LG3), were expressed in early in the development stage (dpp 45 and 75) and reduced expression in late in the development stage (dpp 105, 120 and 135) in both light and dark brown color fruit varieties.

**Table 3:** List of genes detected around the regions of significant SNPs from GWAS result association with the R/B fruit color phenotype. Genes were selected if they have a putatively significant role in fruit ripening and pigmentation. ±150kb on both sides of significant SNP (total 300 Kb) region were considered as a potential region for identifying the possible candidate gene.

| Tag SNP | LG | Gene Name | Distance from Tag SNP<br>(Gene location) |
| --- | --- | --- | --- |
| LG4s65268794 | LG4 | R2R3-MYB transcription factor | 16 Kb |
| LG10s12886617 | LG10 | Ethylene-responsive transcription factor 12 | 116 kb |
| LG10s12886617 | LG10 | RING/U-box superfamily protein | 3 Kb |
| LG10s12886617 | LG10 | Myb family transcription factor family protein | 12 Kb |
| LG3s906369 | LG3 | Protochlorophyllide reductase, chloroplastic | 3.5 Kb |
| LG3s906369 | LG3 | Basic helix-loop-helix (BHLH) DNA-binding superfamily | 83 Kb |

**Figure 5.**
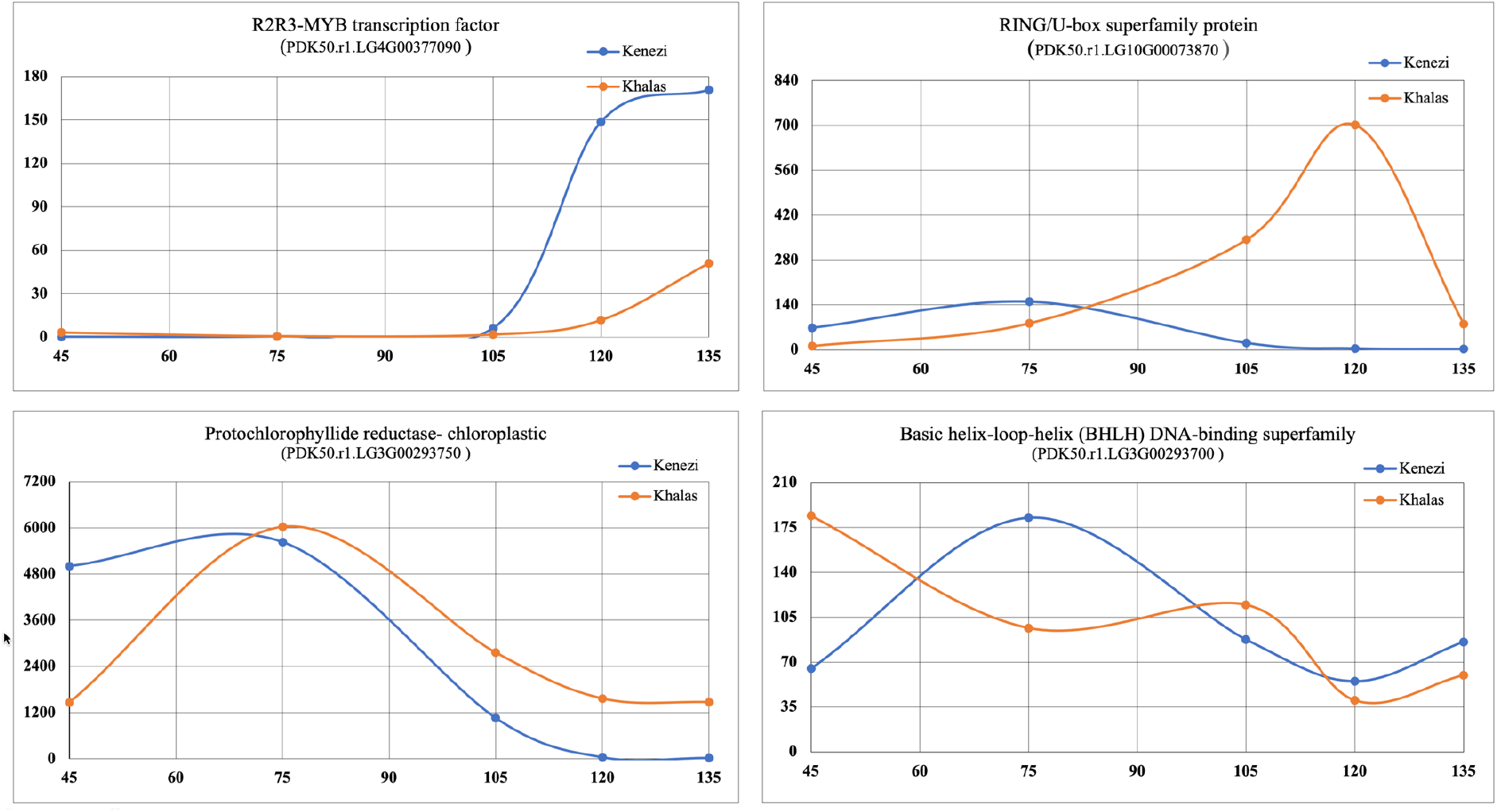
RNA seq analysis of expression of genes related to fruit ripening and pigmentation and are present within the region around identified significant SNPs (Table 2). The gene expression analysis was performed across three or four replicates of fruit development stages of two fruit color varieties, Kenezi (Wild type fruit with red color) and Khalas (VIRIM genotype with yellow color). Each point of the X-axis is the mean of normalized expression read count across three more replicates, and the Y-axis is the time point fruit development stage (post pollination date)

We conducted structural variation analysis on all potential regions from all LGs. The analysis showed a 5kb deletion in the candidate region of SNP LG10s12886617 on LG 10 (Supplementary Figure 12). While this 5kb deletion is located 800 bases away from the pentatricopeptide repeat-containing protein gene and 7Kb away from the SNP LG10s12886617, association of its presence/absence to the GWAS SNP was not high enough to warrant further investigation. The SNPs and INDELs from all potential regions were annotated using SNPEff software and filtered based on LD R2 value >=0.6 and putative impacts value High, Moderate and Modifier. The SNP’s filter results show that a total of 34 SNPs are present across the potential region, 12 non-synonymous variants, one frameshift variants, and 21 three-prime and five prime UTR variants (Supplementary file 4). The SNP list has only one SNP (LG10s12771512) from the gene list mentioned in Table 2. SNP LG10s12771512 is within the Ethylene-responsive transcription factor12 gene from LG10. SIFT results of amino acid substitution effects on protein function analysis showed that the SNP is putatively deleterious possibly affecting protein function.

## Discussion

This study aimed to identify loci and possible candidate genes associated with the color phenotype of tamar stage (dry) date fruit. In our association study, we used genotypic and image data of 188 QC filtered dry date fruit samples. Using the FarmCPU GWAS method and R/B color phenotype, we identified six significant SNPs from the GWAS result (based on the FDR adjusted p-value cut-off) associated with the color phenotype (Table 2). These SNPs span over linkage groups LG3, LG4, LG5, LG10 and one unplaced scaffold MU008982. Among these, the highly significant SNP LG4s65268794 is linked to the previously discovered VIR gene responsible for yellow or red fresh fruit color. The remaining SNPs (five as new loci from this study) are also significantly associated (Wilcoxon statistical test) with fruit color though likely either only for the dry fruit stage or finer color detail in fresh fruit(figure 3a). Possible candidate genes were mapped from the regions surrounding significant SNPs. As expected from previous studies (K. M. Hazzouri et al., 2015; Khaled M. Hazzouri et al., 2019), the R2R3-MYB transcription factor gene is present on LG4 and located 16kb away from the significant SNP LG4s65268794. We confirmed that the this previously identified VIR genotype also has a major effect even on dry fruit color in our samples likely stemming from the starting color at the fresh fruit stage of yellow or red (Supplementary Figure 10).

Beyond the fruit color classification provided by the genotype of the R2R3-MYB transcription factor gene, we investigated how the genotypic variation of other SNPs (Table 2) associated with the color phenotype of dry fruit. That is, do these SNPs provide further genetic contribution to the dry date fruit color phenotype? To do that end we assessed our SNP associations within the homozygous VIR wild type and homozygous VIRIM groups. The results revealed that the newly identified SNPs, excluding SNP LG4s19036701, could give further resolution to the light and dark color fruit on top of fruit color classification by the VIR genotype (Figure 3b & 3c). For some SNPs we think that low numbers of samples in the two groups (wild type group=25 and VIRIM group=61) might be the reason for the lack of the statistically significant association. However, the overall picture is that the newly identified loci are associated with the color phenotype and can distinguish the dry fruit color beyond simply the color observed in fresh fruit. This association may relate to genetic control during the fruit ripening process.

Based on literature search and Blast2GO gene ontology analysis, 6 genes were identified in the regions surrounding significant SNPs (Table 2) that play a role in fruit ripening and pigmentation in other plants (Table 3). RNA-Seq analysis reveals that many of these genes are differentially expressed early or late in the development stages of both light (Khalas) and dark (Kenezi) color fruit (Figure 5). The genes identified, such as the Ethylene-responsive transcription factor12 gene and RING U-Box superfamily protein, are present in the candidate genomic region of SNP LG10s12886617 (LG 10). Ethylene-responsive transcription factor-12 gene is similar to the DORNROSCHEN-like protein in Arabidopsis thaliana. It contains the AP2 domain and has a significant role in the ethylene-activated signalling pathway and cytokinin signalling pathway (Das et al., 2012; Phukan, Jeena, Tripathi, & Shukla, 2017). In Arabidopsis, cytokinin signalling increases the sugar-induced anthocyanin biosynthesis (Das et al., 2012). The candidate region from LG10 also contains an uncharacterised protein (gene id: PDK50.r1.LG10G00073880) which contains Myb DNA-binding 3 domain (Ambawat, Sharma, Yadav, & Yadav, 2013). The Protochlorophyllide reductase gene is present withing the region surrounding LG3s906369 SNP. This gene plays a vital role in chlorophylls’ biosynthetic pathway (Garrone, Archipowa, Zipfel, Hermann, & Dietzek, 2015; Yamazaki, Nomata, & Fujita, 2006).

SNPs that were filtered out due to high FDR, yet remained near the top of our list also identified regions with many genes related to fruit ripening and pigmentation (Kaler, Gillman, Beissinger, & Purcell, 2020; Y. M. Zhang, Jia, & Dunwell, 2019). We used the unadjusted p-value 10e-7 as a cut-off value for identifying those lists of significant SNPs (Supplementary Table 2) however other SNPs may just be below the threshold of significance based on sample numbers used here. The candidate region of SNP LG13s8766984 (LG13) contains an AP2-like ethylene-responsive transcription factor. Other genes include 4-coumarate-CoA ligase and 4-coumarate:coA ligase 3, Myb family transcription factor family protein, AP2/B3-like transcriptional factor family protein, and Chalcone-flavanone isomerase that are present around the region of SNP LG5s4683788 (LG5). Chalcone Isomerase is a critical enzyme for the anthocyanin biosynthesis (J. H. Kang et al., 2014; Sun et al., 2019). The 4-coumarate: CoA ligase is a key enzyme in phenylpropanoid metabolism in plants (Y. Li, Kim, Pysh, & Chapple, 2015; C. H. Wang et al., 2016). Metabolome study of dates detected countable enrichment of phenylpropanoids in the early development of dates (Diboun et al., 2015). Our gene expression analysis shows 4-coumarate: CoA ligase genes (gene id: PDK50.r1.LG5G00393990 and PDK50.r1.LG5G00393890) are highly expressed in early in dark color fruit compared with light color, peaking at 45-75 days post pollination (Supplementary Figure 13). These genes may provide candidates for further study if larger sample numbers reveal them to be indeed be significantly associated with fruit color.

By combining the genotypic data of extensively diverse samples collected from 14 countries and the color phenotype of dry fruit (tamar stage fruit), we successfully performed a GWAS using the FarmCPU method. We identified multiple significant loci and possible candidate genes associated with the color variation of fruit. The new SNPs association with the color of dry date fruit will help add resolution to our understanding of genetic control of commercially important phenotypes in this fruit crop.

## Supporting information

https://wcmq.box.com/s/gw40nlonvs5hg545fz4btk5x1stv1k3x

