## Supplementary material for "Genome-wide analysis of dry (Tamar) Date palm fruit color": https://wcmq.box.com/s/gw40nlonvs5hg545fz4btk5x1stv1k3x

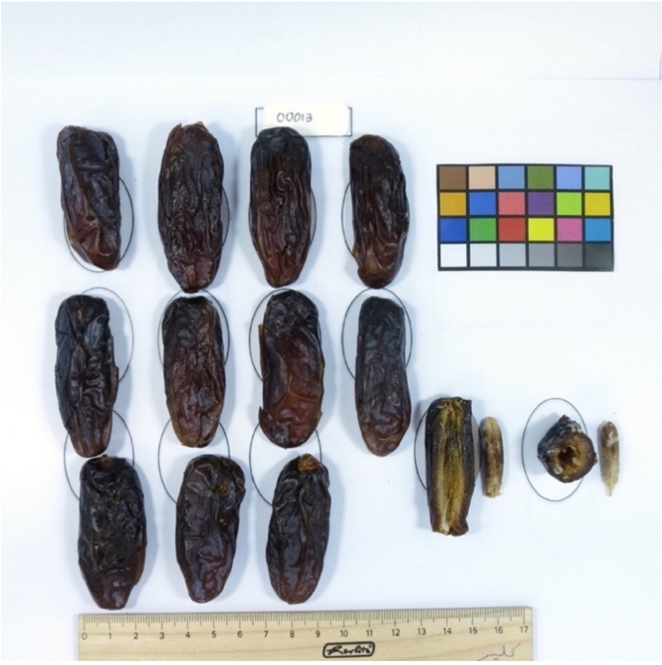


**Supplementary Figure 1**: Fruits Photograph used for measuring the color intensity of fruit color phenotype. The average intensity of each color channel (RGB) from each fruit image was calculated and used as phenotypic data for the association mapping study.

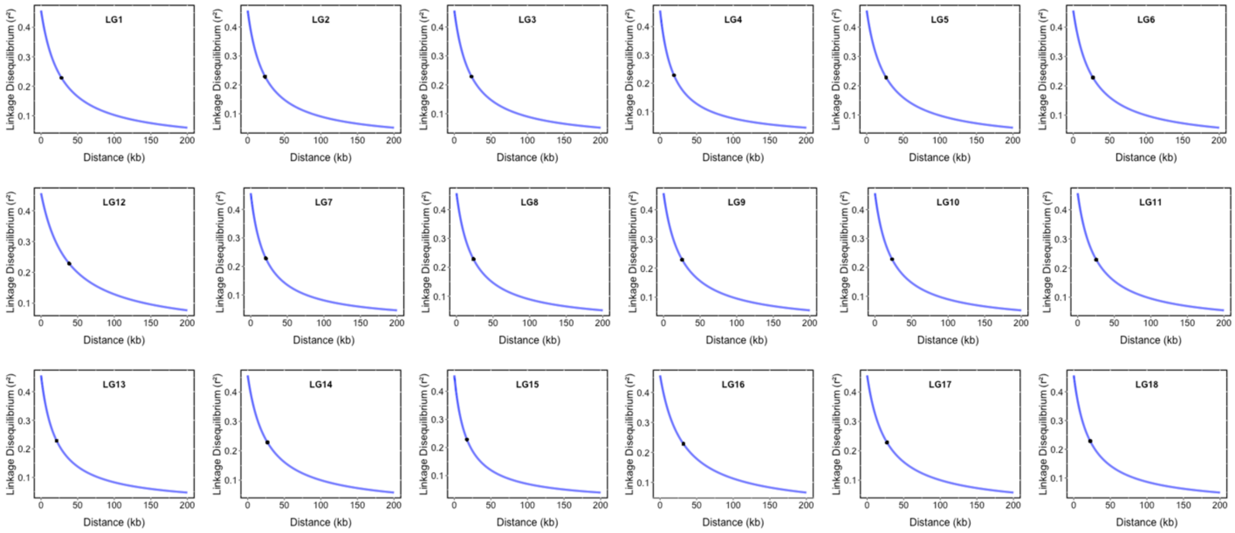


**Supplementary Figure 2:** Decay of Linkage Disequilibrium (LD) with physical distance in each Linkage group (LG). Points on the decay curve represent the half decay distance for each LG.

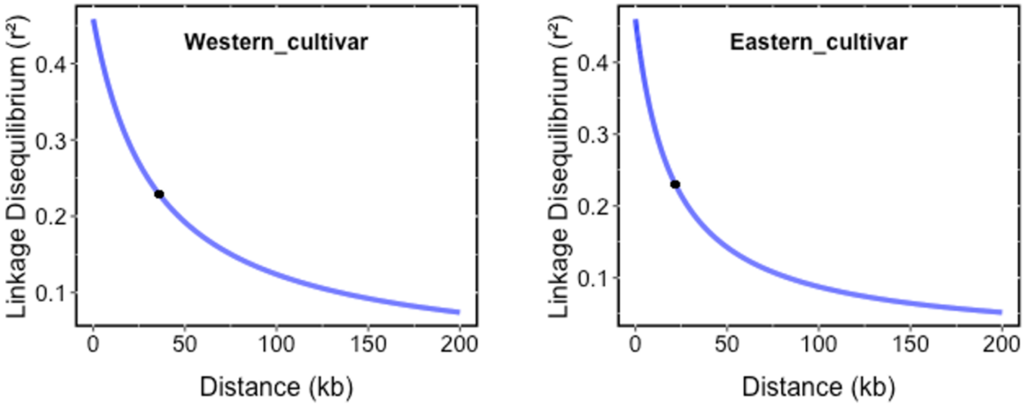


**Supplementary Figure 3:** Decay of Linkage Disequilibrium  (LD) with physical distance in Western and Eastern cultivars. The GWAS mapping population (n=188) contain 68 eastern samples and 106 western samples. The decay of LD with physical distance was calculated in (**a**) Western and (**b**) eastern populations separate. Points on the decay curve represent the half decay distance at 35.85 kb and 21.73 kb for western and eastern populations.

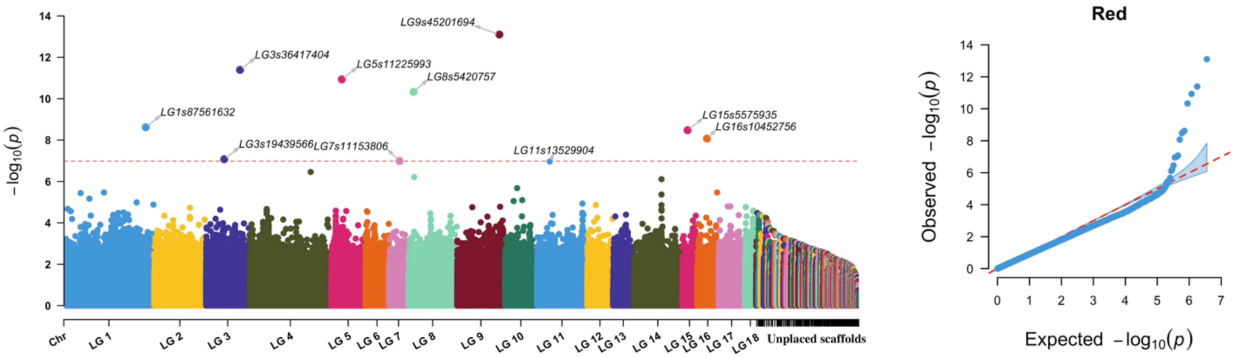


**Supplementary Figure 4:**  Association of Red color channel phenotype of Date palm. LD pruned SNPs set  (3,541,727 SNPs) were used for FarmCPU GWAS.

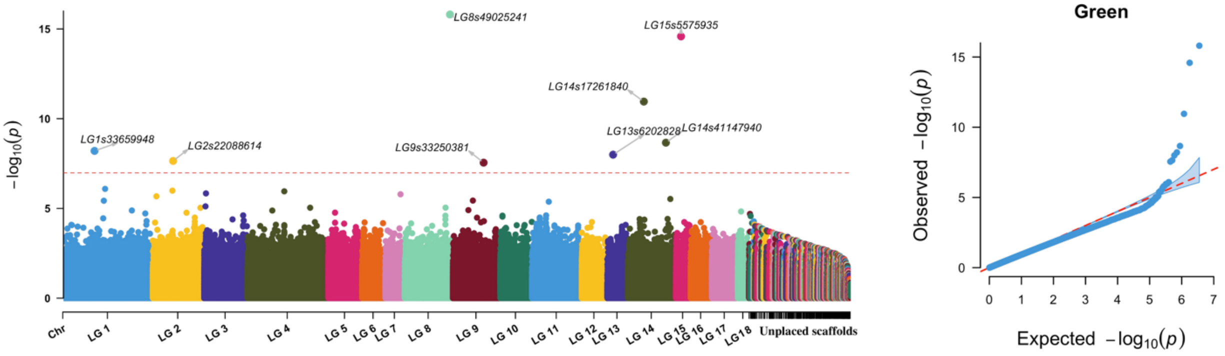


**Supplementary Figure 5:** Association of Green color channel phenotype of Date palm. LD pruned SNPs set (3,541,727 SNPs) were used for FarmCPU GWAS.

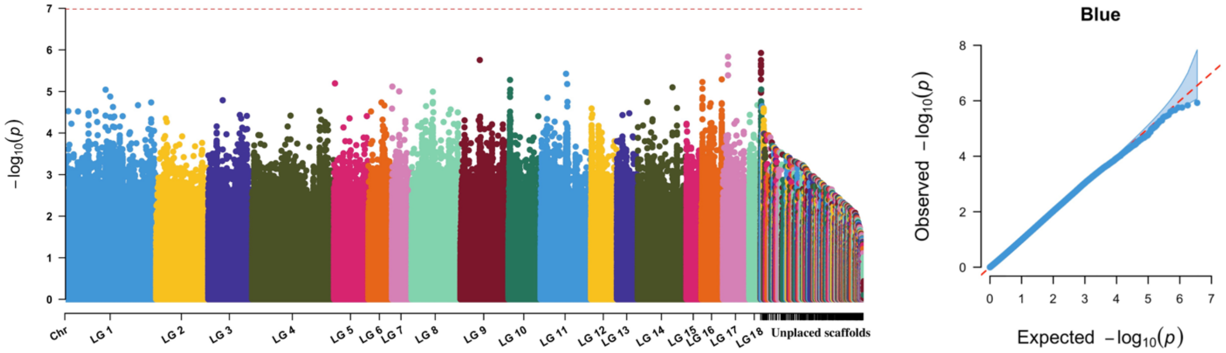


**Supplementary Figure 6 :** Association of Blue color channel phenotype of Date palm. LD pruned SNPs set (3,541,727 SNPs) were used for FarmCPU GWAS.

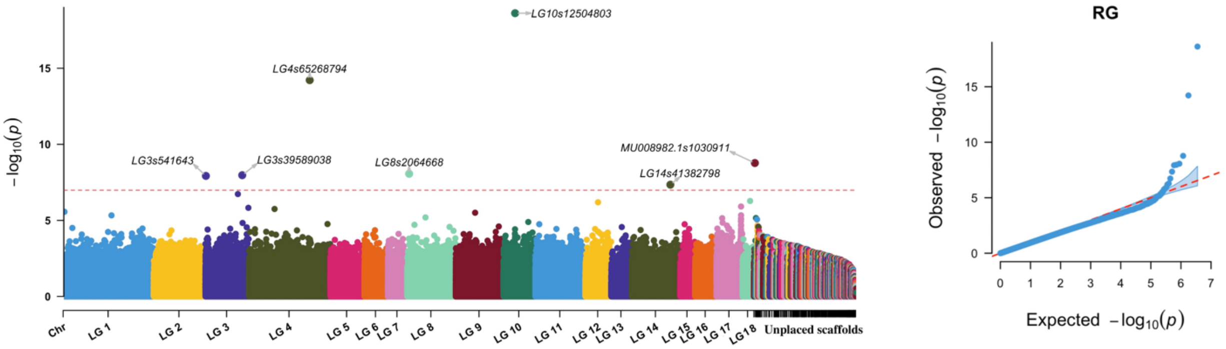


**Supplementary Figure 7:** Association of R/G color channel phenotype of Date palm. LD pruned SNPs set (3,541,727 SNPs) were used for FarmCPU GWAS.

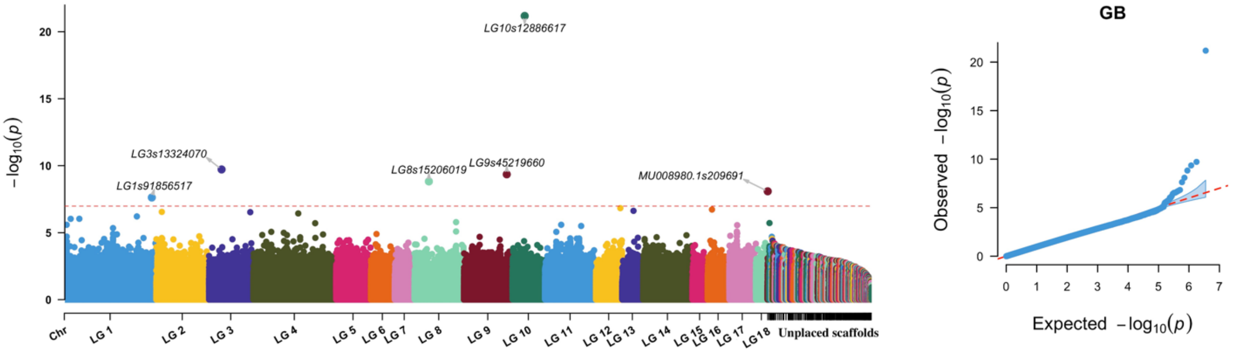


**Supplementary Figure 8:** Association of G/B color channel phenotype of Date palm(n=173). LD pruned SNPs set (3,541,727 SNPs) were used for FarmCPU GWAS.

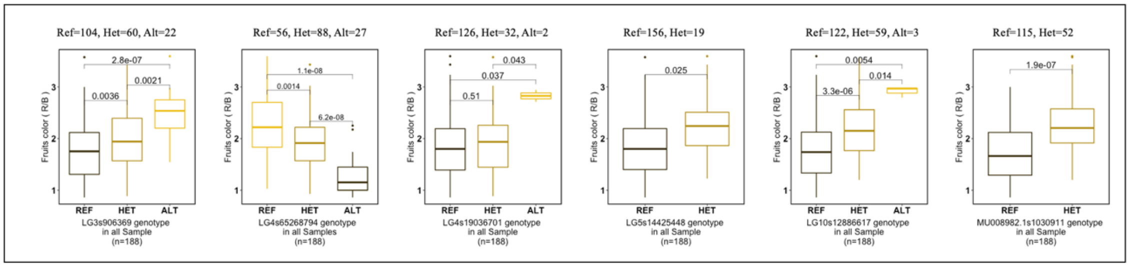


**Supplementary Figure 9:** Boxplot distribution analysis of fruit color phenotype by genotypes of GWAS significant SNPs associated with fruit color (R/B) in all 188 samples. The significant SNPs were identified using FDR adjusted p-value of 0.05 as a cut-off value. Wilcoxon statistical test were used for the P-value calculation.

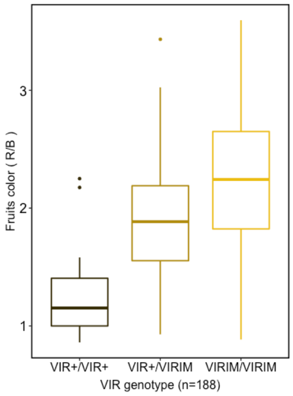


**Supplementary Figure 10**. Boxplot distributions analysis of fruit color phenotypes (R/B) by VIR genotypes showing the VIR genotypic effect on fruit's color. The X-axis represents the VIR genotypes, and Y-axis represents R/B phenotypic value.

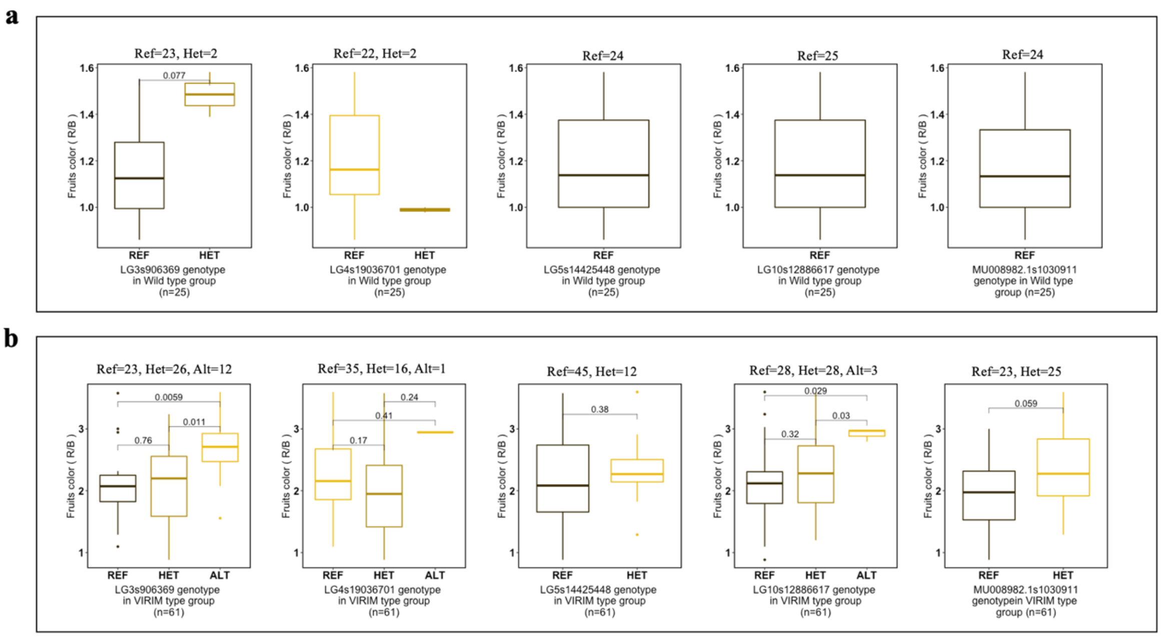


**Supplementary Figure 11**. Boxplot distributions analysis of fruit color phenotype (R/B) by genotypes of significant GWAS SNPs associated with fruit color in VIR sample groups. Samples were classified into two groups: (**a**) wild type group: samples with $VIR^{+}/VIR^{+}$ of $VIR^{+}/VIR^{saf}$ (n=25) and (**b**) VIRIM group: samples with $VIR^{IM}/VIR^{IM}$ genotype (n=61) and then performed the analysis of phenotype by genotypes of significant SNPs. P-values were calculated using by Wilcoxon statistical test. The X-axis represents the SNP's genotypes, and Y-axis represents R/B phenotypic value.

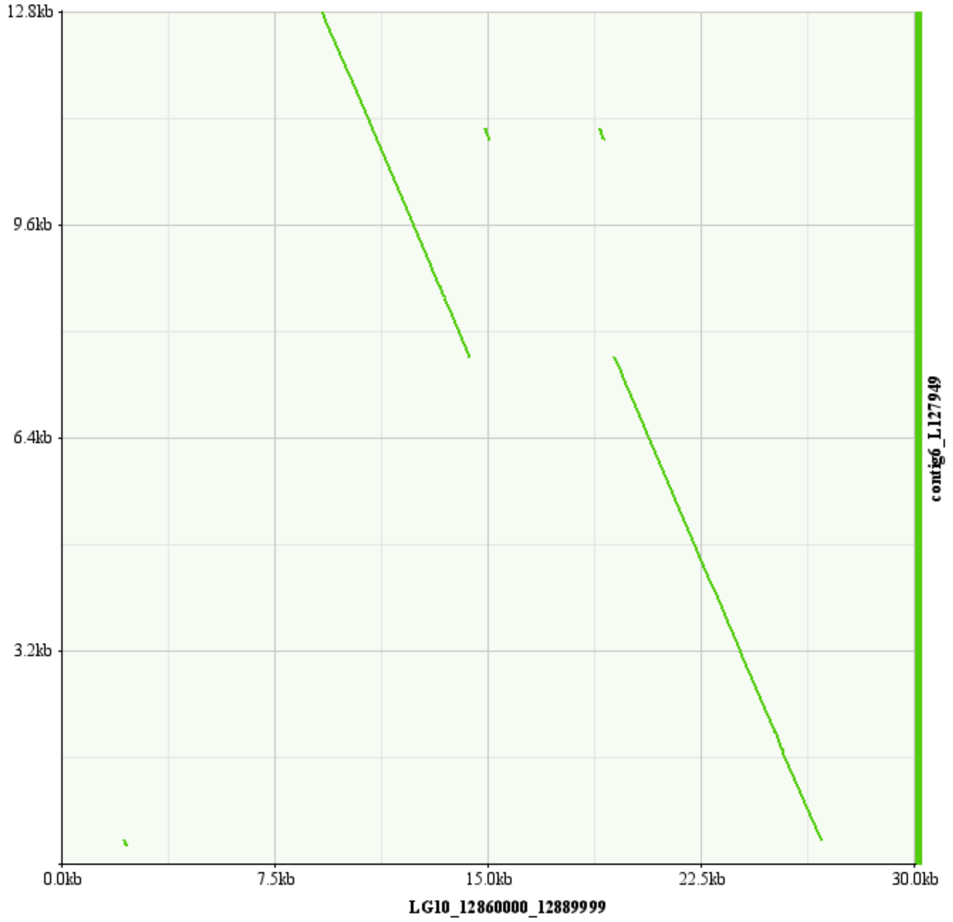


**Supplementary Figure 12:** Structural variation (deletion) present in candidate region of LG10s12886617 SNPs on LG 10 (30 kb region from LG10, 12.86Mb to 12.890Mb, aligned against contig 6). Clipped, discordant, unmapped, and indel reads from samples having LG10s12886617 SNP with homozygous allele alternative (ALT) to the reference genome were assembled and aligned against LG 10. The alignment of contigs 6 (12.7949kb length) showed a 5kb deletion in the region 12.8743Mb to 12.8794Mb of LG 10. This 5kb deletion is located 800 bases from the pentatricopeptide repeat-containing protein gene and 7Kb away from the SNP LG10s12886617.

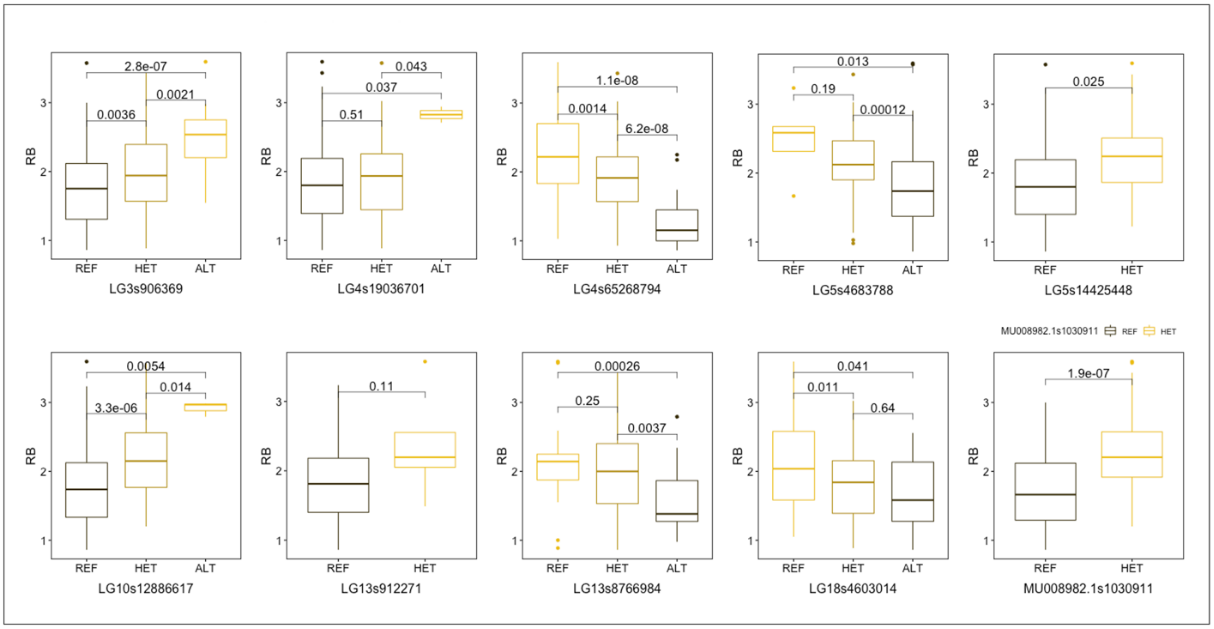


**Supplementary Figure 13**. Boxplot distributions analysis of fruits color phenotype by genotypes of significant GWAS SNPs in all 188 samples. The SNPs were identified using an unadjusted p-value 10e-7 as a cut-off value for identifying the list of significant SNPs associated with the fruit's color phenotype (R/B).

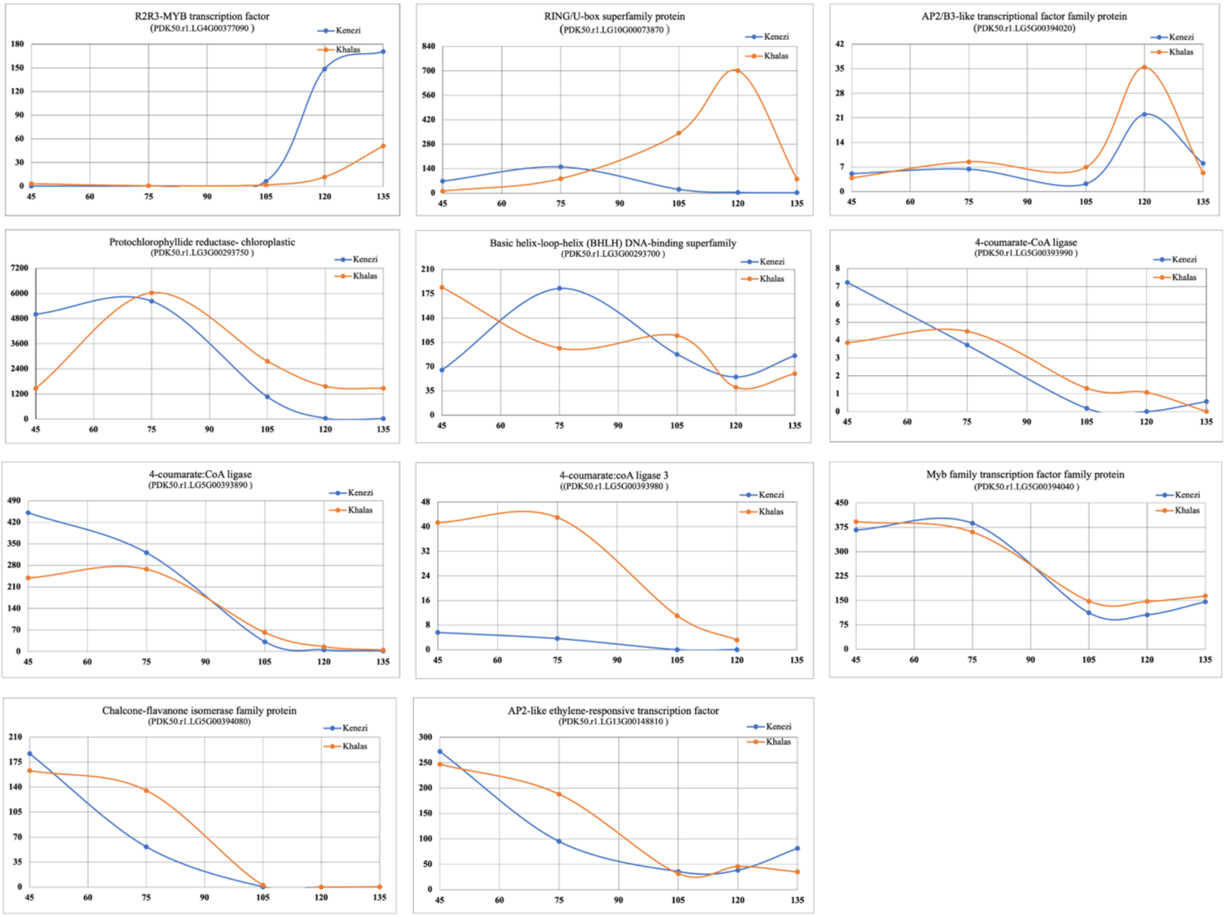


**Supplementary Figure 14**. RNA seq analysis of expression of genes related to fruit ripening and pigmentation are present around the region of significant SNPs identified by using an unadjusted p-value 10e-7 as a cut-off value. The gene expression analysis was performed across three or four replicates of fruit development stages of two fruit color varieties, Kenezi (Wild type fruit with red color) and Khalas (VIRIM genotype with yellow color). Each point of the X-axis is the mean of normalised expression read count across three more replicates, and the Y-axis is the time point fruit development stage (post pollination date)

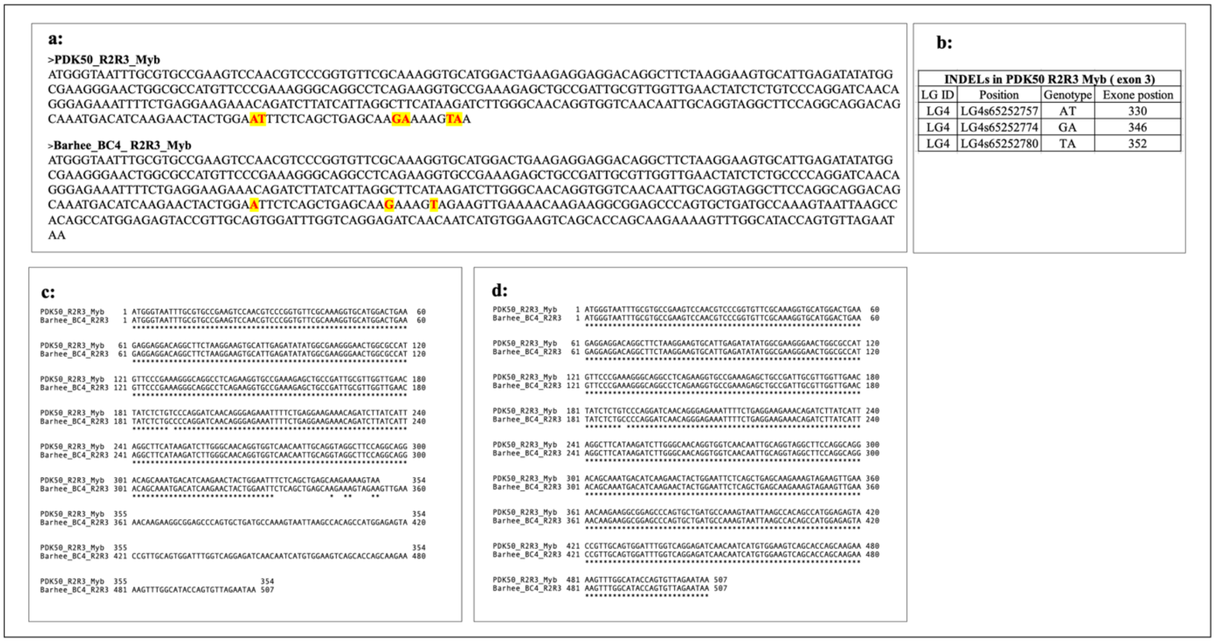


**Supplementary Figure 15:** Correction of Indel mutation in R2R3 Myb transcription factor gene sequence from PDK50 genome. Due to sequencing error, three indel mutations occurred in exon positions 330, 346 & 352 of the R2R3 Myb transcription factor gene. These errors could not be resolved in pilon genome polishing due to less read coverage (minimum reads used for pilon polishing was 10), and it caused a premature stop codon at the 344$^{th}$ exon position of the gene. Due to the premature stop codon, the gene became 354bp sequence length and lost 157 bp. We compared the 354bp length sequence of the R2R3 Myb transcription factor gene along with the missing 157bp sequence against the  R2R3 Myb transcription factor gene from the Barhee BC4 genome and corrected the sequencing errors. The pairwise. alignment shows that the missing 157bp sequence is 100% similarity to the R2R3 Myb transcription factor gene from the Barhee BC4. **a :** R2R3 Myb transcription factor gene sequence from PDK50 genome (354bp sequence length) and Barhee BC4 genome (507bp sequence length) . Indel mutations were marked in red color. **b:** Indel mutations details of R2R3 Myb transcription factor gene. **c:** pairwise alignment of 354bp sequence of R2R3 Myb transcription factor gene against R2R3 Myb transcription factor gene from Barhee. **d:** Alignment of error corrected sequence of R2R3 Myb transcription factor gene including the missing 157bp sequence against R2R3 Myb transcription factor gene from Barhee BC4 genome.

**Supplementary Table 1:** Genomic coordinates of VIR *gene,* retrotransposon insertion (Ibn Majid), and locations of start lost codon and termination codon variant in LG 4 of PDK50 genomes.

| Allele TYPE | Features | Start | End |
|---|---|---|---|
| Retrotransposon Insertion (VIRIM) | Exon1 | 65251488 | 65251623 |
| | Exon2 | 65251695 | 65251824 |
| | Exon3 | 65252695 | 65252935 |
| | Ibn Majid (LTR retrotransposon) | 65252928 | 65264522 |
| | Termination codon | 65252936 | 65252938 |
| Lost start codon (VIRsaf) | ATG/ATA SNP | 65251490 | 65251490 |
| Wild Type ( VIR+) | Exon 3 (CDS), 5' of Ibn Majid | 65252695 | 65252927 |
| | Termination codon (TGG/TGA) | 65264705 | 65264707 |

**Supplementary Table 2**: List of significant SNPs significantly associated with the (R/B) phenotype. An unadjusted p-value of 10e-7 was used as a cut-off value for identifying the list of significant SNPs.

| **SNP** | **CHR** | **Position** | **P.value** | **maf** | **FDR P.values** |
|---|---|---|---|---|---|
| LG4s65268794 | LG4 | 65268794 | 7.40E-12 | 0.416201117 | 2.62E-05 |
| LG10s12886617 | LG10 | 12886617 | 1.56E-10 | 0.19273743 | 0.000276496 |
| LG3s906369 | LG3 | 906369 | 3.29E-10 | 0.293296089 | 0.000388953 |
| LG4s19036701 | LG4 | 19036701 | 3.84E-09 | 0.17877095 | 0.003401362 |
| LG5s14425448 | LG5 | 14425448 | 4.98E-08 | 0.089385475 | 0.035277173 |
| MU008982.1s1030911 | MU008982.1 | 1030911 | 7.25E-08 | 0.203910615 | 0.042803883 |
| LG13s8766984 | LG13 | 8766984 | 2.58E-07 | 0.446927374 | 0.130612832 |
| LG5s4683788 | LG5 | 4683788 | 6.45E-07 | 0.170391061 | 0.26190628 |
| LG18s4603014 | LG18 | 4603014 | 6.66E-07 | 0.301675978 | 0.26190628 |
| LG13s912271 | LG13 | 912271 | 7.89E-07 | 0.11452514 | 0.279481236 |

**Supplementary Table 3**: List of genes detected around the potential region of SNPs from GWAS results associated with R/B phenotypes to have significant role in fruit ripening and pigmentation. SNPs were identified based on an unadjusted p-value cut-off of 10e-7.

| **Tag SNPs** | **LG ID** | **Gene ID** | **Gene Description** | **Distance from Tag SNP (gene location)** |
|---|---|---|---|---|
| LG3s906369 | LG3 | PDK50.r1.LG3G00293750 | Protochlorophyllide reductase, chloroplastic | 3.5 Kb |
| LG3s906369 | LG3 | PDK50.r1.LG3G00293700 | Basic helix-loop-helix (BHLH) DNA-binding superfamily | 83 Kb |
| LG4s65268794 | LG4 | PDK50.r1.LG4G00377090 | R2R3-MYB transcription factor | 16 Kb |
| LG5s4683788 | LG5 | PDK50.r1.LG5G00393890<br>PDK50.r1.LG5G00393990 | 4-coumarate:CoA ligase | 133 Kb |
| LG5s4683788 | LG5 | PDK50.r1.LG5G00393910.1<br>PDK50.r1.LG5G00393950.1 | AP2/ERF and B3 domain-containing transcription factor family | 88 Kb |
| LG5s4683788 | LG5 | PDK50.r1.LG5G00393980 | 4-coumarate:coA ligase 3 | 50 Kb |
| LG5s4683788 | LG5 | PDK50.r1.LG5G00394020 | AP2/B3-like transcriptional factor family protein | 8 Kb |
| LG5s4683788 | LG5 | PDK50.r1.LG5G00394040 | Myb family transcription factor family protein | 12 Kb |
| LG5s4683788 | LG5 | PDK50.r1.LG5G00394080 | Chalcone-flavanone isomerase family protein | 87 kb |
| LG10s12886617 | LG10 | PDK50.r1.LG10G00073810 | Ethylene-responsive transcription factor 12 | 116 Kb |
| LG10s12886617 | LG10 | PDK50.r1.LG10G00073870 | RING/U-box superfamily protein | 3 Kb |
| LG10s12886617 | LG10 | PDK50.r1.CM021349.1G000 73880.1 | Myb family transcription factor family protein | 12 kb |
| LG13s912271 | LG13 | PDK50.r1.LG3G00148810 | AP2-like ethylene-responsive transcription factor | 36Kb |
